# Combinatorial optimization of pathway, process and media for the production of p-coumaric acid by *Saccharomyces cerevisiae*

**DOI:** 10.1101/2023.11.28.568970

**Authors:** Sara Moreno-Paz, Rianne van der Hoek, Elif Eliana, Vitor A.P. Martins dos Santos, Joep Schmitz, Maria Suarez-Diez

**Affiliations:** Laboratory of Systems and Synthetic Biology, Wageningen University & Research, 6708 WE Wageningen, The Netherlands; Department of Science and Research - dsm-firmenich, Science & Research, 2600 MA Delft, The Netherlands; Bioprocess Engineering Group, Wageningen University & Research, Wageningen 6700 AA, The Netherlands

## Abstract

Microbial cell factories are instrumental in transitioning towards a sustainable bio-based economy, offering alternatives to conventional chemical processes. However, fulfilling their potential requires simultaneous screening for optimal media composition, process and genetic factors, acknowledging the complex interplay between the organism’s genotype and its environment. This study employs statistical Design of Experiments (DoE) to systematically explore these relationships and optimize the production of p-coumaric acid (pCA) in *Saccharomyces cerevisiae*. Two rounds of fractional factorial designs were used to identify factors with a significant effect on pCA production, which resulted on a 168-fold improvement on pCA titer. Moreover, a significant interaction between the culture temperature and expression of ARO4 highlighted the importance of simultaneous process and strain optimization. The presented approach leverages the strengths of experimental design and statistical analysis and could be systematically applied during strain and bio-process design efforts to unlock the full potential of microbial cell factories.

## 1 Introduction

Microbial cell factories play a pivotal role in driving the transition towards a bio-based economy, being a sustainable alternative to traditional chemical processes [1]. Microorganisms can efficiently transform raw materials into valuable products. However, to unlock their potential for biotransformation in an economically feasible manner, it is essential to optimize production pathways and bio-processes [2].

Pathways can be optimized sequentially by tuning individual genetic factors in isolation. However, this does not capture the complex interplay between different genetic elements and the products they code for. It is hence desirable to perform combinatorial pathway optimization, which is based on the simultaneous optimization of multiple genetic factors and facilitates the identification of complex interactions [3, 4]. Moreover, the overall performance of the microbial cell factory is not only determined by its genotype, but it is also influenced by the production conditions, as factors such as media nutrients, pH, cultivation temperature and aeration influence cell physiology and metabolism. Strains are usually optimized holding the environmental conditions constant and only the most promising strain advances to the bio-process optimization stage [5, 6]. However, this approach might ignore genetic designs that, although inferior in standard laboratory conditions, have bigger potential when the media and bio-process are optimized [7]. Only by simultaneously screening for optimal media composition and genetic factors, the dynamic interplay between the organism’s genotype and the environment in which it operates can be considered [7, 8].

Combinatorial optimization of strains, media and process parameters, however, requires exponentially increasing resources. Statistical design of experiments (DoE) allows a structured exploration of the relationships between experimental variables (factors) and the measured response. Full factorial designs are a type of DoE designs that test all possible combinations of factor levels, characterizing factor effects and allowing the estimation of interactions. The number of experiments to be performed depends on the number of genetic and environmental factors to be tested (e.g. expression of a gene, temperature) and the number of levels per factor (e.g. low, medium and strong gene expression, 20°C, 25°C, 20°C and 35°C) according to 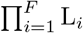, where *F* is the number of factors and *L*_*i*_ is the number of levels of factor *i*. In these designs, the effect of a factor is not only estimated considering replicate experiments, but also all the experiments where the given factor is constant regardless of other factor’s levels. This property can be leveraged in fractional factorial designs that reduce the number of experiments to perform while maximizing the information gain. This is achieved by performing experiments that preserve orthogonality in the desired factors, i.e. ensuring that the effect of a factor is not confounded by planned changes in other factors. The generated data is fitted to a linear model so main effects (MEs), representing the impact of not-confounded factors on the response, are identified. Similarly, the so-called Two Factor Interactions (2FI) that occur when the effect of a factor on the response changes based on the level of another factor, can also be estimated [9].

We used production of p-coumaric acid (pCA) by *Saccharomyces cerevisiae* as an example of DoE-aided combinatorial pathway, media and process optimization. pCA can be produced from phenylalanine (Phe), an aromatic amino acid produced within the shikimate pathway. It is a precursor for a wide array of biologically relevant molecules such as pharmaceuticals, flavors, fragrances and cosmetics [10]. Although pCA production has been independently optimized at the strain and bio-process levels [10–14], we show the interplay between genetic and environmental factors highlighting the importance of simultaneous process and strain optimization.

## 2 Experimental procedures

### 2.1 Strain construction

Promoter, terminator and ORFs sequences from *aro4, aroL, aro7, pal1, c4h* and *cpr* codon optimized for *S. cerevisiae* were obtained from Moreno-Paz et al. [15] (Sup. Table 1). Twelve cassettes formed by combinations of promoter, ORF and terminator (Sup. Table 1) were assembled via Golden Gate into a backbone plasmid containing a 50 bp homologous connector sequence to facilitate *in vivo* recombination of the gene cluster [16]. Golden Gate products were transformed into chemically competent *E. coli* DH10B cells, plasmids were isolated and cassettes were confirmed by PCR as described in Moreno-Paz et. al [15].

Strains were constructed as described in Moreno-Paz et al [15]. In short, a host strain with Cas9 integrated in the non-coding region between YOR071c and YOR070c in chromosome 15 was transformed with a linear guide RNA targeting AEHG01000256.1 (210ng/kb), equimolar cassettes for the required designs (Table 1, Sup. Table 2) (100-300ng/kb) and linear backbone fragments (35ng/kb) following the LiAc/ssDNA/PEG method [17]. The connector sequences on the cassettes facilitates *in vivo* recombination of a cluster of genes in the genome [16]. Transformants were plated on Qtray (NUNC) with 48-divider (Genetix) containing YEPhD agar medium and selection agent. Colonies appeared on plate after 3 days of incubation at 30°C. Single colonies were picked with Qpix 420 (Molecular Devices) into 96 well plate containing YEPhD agar medium and selection agent and regrown for 3 days at 30°C. Colonies were confirmed using whole genome sequencing as described in Moreno-Paz et al. and correct strains were stored in 10% glycerol at -80 °C [15].

**Table 1:** Structure of the gene clusters. Cells values indicate the promoter used for each gene. Promoters were selected from **ML paper**.

| Gene cluster | ARO4 | AROL | ARO7 | PAL1 | C4H | CPR1 |
| --- | --- | --- | --- | --- | --- | --- |
| 1 | TDH3 | TEF1 | ACT1 | RPS9A | CHO1 | CCW12 |
| 2 | TDH3 | TEF1 | ACT1 | VMA6 | PXR1 | CCW12 |
| 3 | TDH3 | TEF1 | PFY1 | RPS9A | CHO1 | CCW12 |
| 4 | TDH3 | TEF1 | PFY1 | VMA6 | PXR1 | CCW12 |
| 5 | TDH3 | RPL28 | ACT1 | RPS9A | CHO1 | CCW12 |
| 6 | TDH3 | RPL28 | ACT1 | VMA6 | PXR1 | CCW12 |
| 7 | TDH3 | RPL28 | PFY1 | RPS9A | CHO1 | CCW12 |
| 8 | TDH3 | RPL28 | PFY1 | VMA6 | PXR1 | CCW12 |
| 9 | MYO4 | TEF1 | ACT1 | RPS9A | CHO1 | CCW12 |
| 10 | MYO4 | TEF1 | ACT1 | VMA6 | PXR1 | CCW12 |
| 11 | MYO4 | TEF1 | PFY1 | RPS9A | CHO1 | CCW12 |
| 12 | MYO4 | TEF1 | PFY1 | VMA6 | PXR1 | CCW12 |
| 13 | MYO4 | RPL28 | ACT1 | RPS9A | CHO1 | CCW12 |
| 14 | MYO4 | RPL28 | ACT1 | VMA6 | PXR1 | CCW12 |
| 15 | MYO4 | RPL28 | PFY1 | RPS9A | CHO1 | CCW12 |
| 16 | MYO4 | RPL28 | PFY1 | VMA6 | PXR1 | CCW12 |

### 2.2 pCA production experiments

Single colonies were grown in 10 ml YPDA media (Takara) in 50 ml tubes for 24h. Cultures were washed and inoculated in minimal media at starting ODs of 0.3 or 0.6 according to the experimental design. Minimal media contained 20 g/l glucose (Acros Organics), and 1.7 g/l yeast nitrogen base without amino acids or ammonium sulfate (BD Difco). A 60.5 mM nitrogen concentration in the media was obtained with 4g/l ammonium sulfate (Acros Organics) or 1.82 g/l urea (Acros Organics). When required, media was buffered at a pH of 7 using 126 mM Na_2_HPO_4_ (Acros Organics) and 18 mM citric acid (Sigma-Aldrich) [18] and/or supplemented with 5 mM Phe (Sigma-Aldrich) and/or 5 mM Glu (Sigma-Aldrich). Cells were grown for 48 h at the required temperature and agitation speed in 50 ml mini-bioreactor tubes (Corning) in an Innova 44 incubator (New Brunswick Scientific). At the end of the cultivation samples for OD measurements and pCA quantification were taken.

### 2.3 pCA quantification

For pCA quantification, 400 μl of culture were mixed with 800 μl acetonitrile (Thermo Scientific) and centrifuged for 10 min at 4000 g. The acetonitrile phase was used for analysis using high performance liquid chromotagoraphy (HPLC) on a Shimadzu LC2030C Plus 2 machine equipped with a Poroshell 120EC-C18 column (250 x 4.6 mm, Agilent) and a UV/vis detector. Mobile phase was used at a rate of 1 ml/min and was composed of Milli-Q water (A), 100 mM formic acid (B), and acetonitrile (C) at varying proportions. The concentration of B was fixed at 10% and the concentration of C varied from 13.5% to 67.5% during the first 12 min, was kept constant at 67.5% for 5 min, decreased to 13.5% in 2 min and kept this concentration for 6 min. pCA was detected at a wavelength of 280 nm. Standards were prepared using pCA purchased from Sigma-Aldrich.

### 2.4 Experimental design and statistical analysis

The `FrF2` function from the `FrF2` R package was used for the generation of the designs given the number of factors and the desired resolution [19].

Experimental data was used to train a linear model:

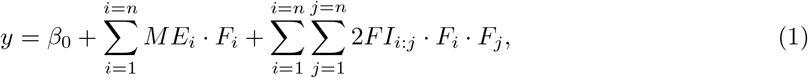

where *y* represents the pCA concentration; *ME*_*i*_ refers to the main effect of factor *i* (*F*_*i*_) and 2*FI*_*i*:*j*_ refers to the two-factor interaction between factor *i* and *j. n* indicates the total number of factors.

Ordinary least squares regression minimizing the sum of squared differences between the observed and predicted values was used to estimate the coefficients for each term in the model (*ME*_*i*_, 2*FI*_*i*:*j*_) using the R `lm` function. Then the `summary` function was used to obtain the ANOVA table which provides the estimated coefficients and their associated p-values. p-Values were corrected using Bonferroni. The adjusted coefficient of determination (R^2^) was used to assess the model fit to experimental data.

## 3 Results

### 3.1 Selection of genetic and environmental factors and levels

The shikimate pathway is tightly regulated and aromatic amino acids exert feedback inhibition on some of its enzymes (Figure 1) [20]. Expression of feedback-resistant variants of ARO4 (ARO4^K229L^) and ARO7 (ARO7^G141S^) are common strategies to increase pCA production [10, 12]. Besides, the phosphorylation of shikimate performed by ARO1 has been hypothesized as rate limiting step. Rodriguez et al. reported a beneficial effect of expressing *E. coli* AROL to increase the flux through this reaction [12]. Therefore, we selected the expression of ARO4^K229L^, ARO7^G141S^ and AROL as genetic factors with the potential to affect pCA production. Each of these factors was evaluated at two levels based on the strength of the promoter-terminator pair assigned to each gene (Table 2) [15].

**Figure 1:**
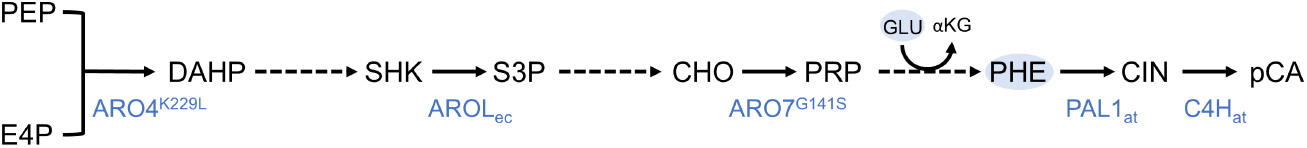
pCA production pathway. Genes which expression is considered as factor for the design are shown. Glu and Phe are highlighted as they are selected as factors for media optimization. PEP, phosphoenolpyruvate; E4P, erithrose-4-phosphate; DAHP, 3-deoxy-7-phosphoheptulonate; SHK, shikimate; S3P, shikimate-3-phosphate; CHO, chorismate; PRP, phrephenate; PHE, phenylalanine, CIN, cinnamate; pCA, p-coumaric acid; GLU, glutamate; αKG, α-ketoglutarate.

**Table 2:** Factors and levels used for pCA optimization.

| Factor | Low-level (-1) | High-level (1) |
| --- | --- | --- |
| Temperature | 20 °C | 30 °C |
| Agitation | 180 rpm | 250 rpm |
| Initial cell density | 0.3 | 0.6 |
| N-source | Urea | Ammonium sulfate |
| pH | Unbuffered | Buffered |
| Phe | 0 | 5 mM |
| Glu | 0 | 5 mM |
| PAL-C4H promoters | VMA6 - PXR1 | RPS9A - CHO1 |
| ARO7 promoter | PFY1 | ACT1 |
| AROL promoter | RPL28 | TEF1 |
| ARO4 promoter | MYO4 | TDH3 |

To produce pCA from Phe, the expression of two heterologous genes is required: phenylalanine ammonia lyase (PAL) and cinnamate 4-hydroxylase (C4H). Although *S. cerevisiae* contains endogenous cytochrome P450 reductases (CPR), expression of a C4H associated CPR is recommended [10–13]. We expressed *Arabidopsis thaliana* CPR under a constitutive promoter and considered the expression of PAL and C4H as an additional genetic factor for the design. The expression levels of PAL and C4H were evaluated using two promoter-terminator pairs (Table 2).

Temperature (T), agitation (rpm) and initial cell density (OD) are usual variables tuned during bio-process optimization and were selected as factors to improve pCA titers [21–26]. Temperature was varied between 30°C, the optimal growth temperature of *S. cerevisiae*, and 20°C, as lower temperatures might improve heterologous pathway expression (Table 2) [21]. Agitation and initial OD were varied between 180-250 rpm and 0.3-0.6, respectively (Table 2).

Combes et al. showed that the pH of the media affects pCA production [27]. Acidic pH, below pCA pKA (4.65), favours the undissociated form of pCA (pHCA) in the media that diffuses into the cell where it dissociates (pCA^-^), acidifying the cytoplasm and requiring active export at the cost of ATP. Considering this, two factors that influence the pH of the media were selected: the addition of a buffer and the use of different nitrogen sources (Table 2). When ammonium sulfate or urea are used as nitrogen source, pH below and above pCA pKA are expected respectively [18]. Independently of the N-source used, the citrate phosphate buffer can control the pH of the culture but can negatively impact cell growth [18].

Media supplementation is an additional common strategy to increase production [6, 26, 28, 29]. Phenylalanine is the substrate of PAL, the first enzyme required for the production of pCA and glutamate is the nitrogen donor used during Phe production (Figure 1) . Therefore, the additions of these amino acids were considered as additional factors (Table 2).

### 3.2 Resolution IV design: impact of individual factors on pCA production

The effect of changing process conditions and media-related factors on pCA production was evaluated using a resolution IV fractional factorial design. These designs allow the estimation of main effects of all the factors while confounding 2-factor interactions. They can be used during screening to identify factors with a significant impact on production that can be the focus of later optimization.

In order to obtain a resolution IV design with 11 factors (Table 2), 32 experiments are required (Figure 2, Sup. Table 3). Although traditional applications of DoE use single-replicate screening, replicates are a necessity to assess biological variation and, for each experiment, pCA production was measured in three independent cultures [4]. These experiments involved the construction of 16 strains including all possible combinations of the four selected genetic factors. Each strain is then tested in two different conditions determined by the design. However, strains containing gene clusters 2 and 6 (Table 1) could not be constructed and the effect of not performing four out of the 32 experiments was evaluated. Reducing the number of experiments to 28 did not affect the estimation of main effects but increased the complexity of the confounding patterns for the 2FI. Considering the goal of the experiment was to determine the ME, construction of the two additional strains was not required, which accelerated the implementation of the design round. In the 28 experiments performed, pCA production varied two orders of magnitude, from 1.3 mg/l to 158.2 mg/l, confirming the impact of the selected factors on production (Figure 2A).

**Figure 2:**
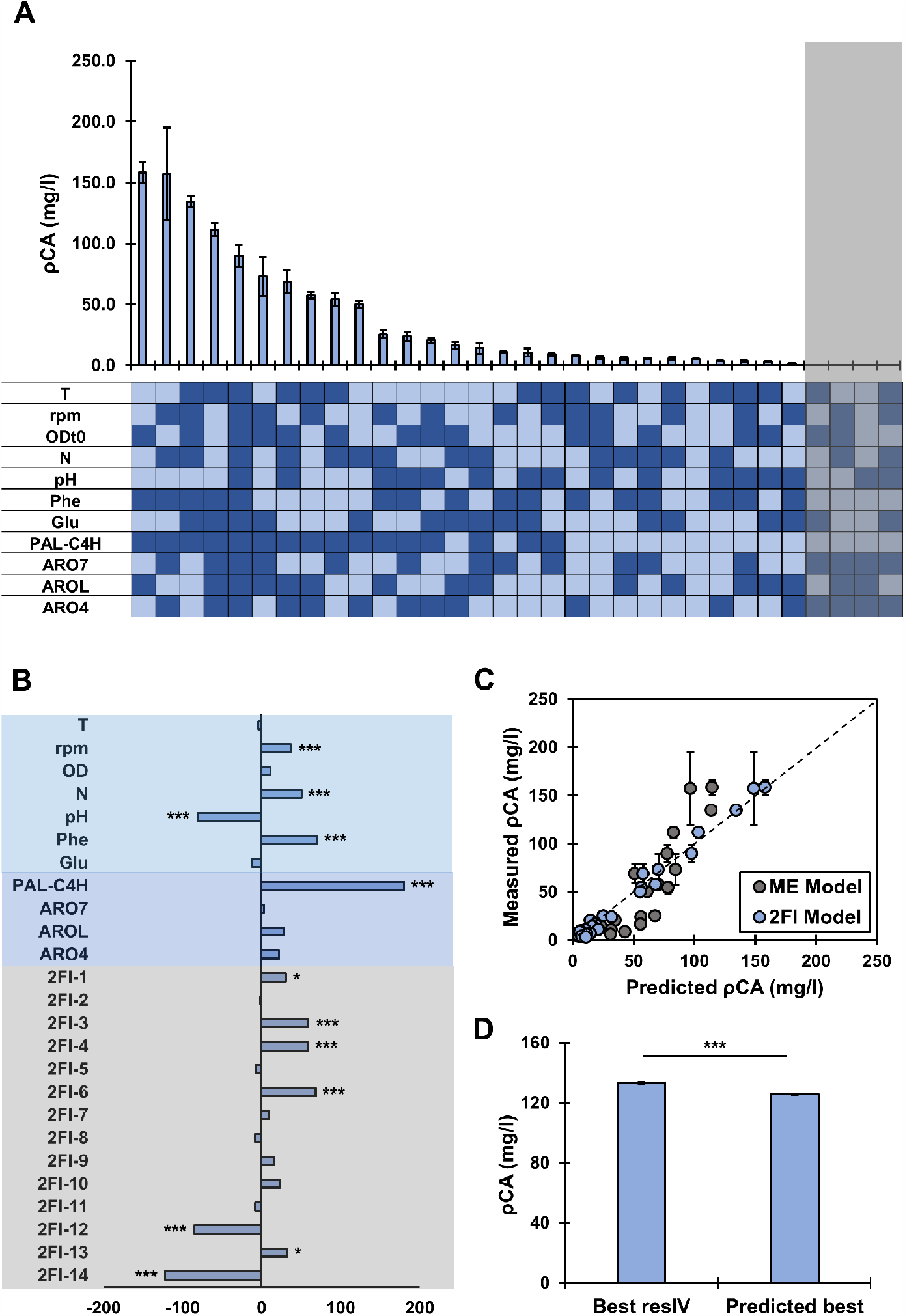
Resolution IV Design **A**. Measured pCA production at different combinations of factors and levels. Light colors indicate low levels (-1) and dark blue indicates high level (1). The shaded gray area indicates the four experiments that could not be performed. **B**. Coefficients of the 2FI model. *** indicates corrected p-value ≤ 0.001, ** ≤ 0.01 and *≤ 0.05. **C**. Fit of models including main effects (ME) or main effects and two-factor interactions (2FI model) to experimental data. **D** pCA production in validation experiment.

Linear models containing only ME or ME and confounded 2FI were trained. Including 2FI increased the coefficient of determination from 0.66 to 0.94 (Figure 2C). Figure 2B shows the estimated coefficients of the model including 2FI. An ANOVA was used to determine the significance of each ME and 2FI on pCA production and p-values were corrected using Bonferroni. All factors but T, OD, Glu and ARO7 had a significant effect on pCA production. Moreover, seven of the estimated 2FI were also significant.

The MEs with the highest impact on performance was the expression strength of PAL and C4H, with a positive regression coefficient. This indicates that a high expression of the heterologous genes for pCA production is essential to obtain high titers. The effect of PAL-C4H was followed by the negative impact of buffering of the media represented by a negative regression coefficient. Although the addition of a buffer could control the dissociation of pCA [27], it negatively affected cell growth and resulted in overall low pCA titers. The third most relevant ME was the addition of Phe, with a positive coefficient that shows the benefit of Phe supplementation on pCA production [30].

Notably, estimated coefficients for 2FI-6, 12 and 14 had a similar impact on pCA titer than PALC4H, pH and Phe and the coefficient of determination was significantly improved when 2FI were considered for pCA production (Figure 2 B, C), indicating that correctly estimating 2FI is required to optimize pCA production. The importance of 2FI was confirmed in an independent experiment where the best experiment from the resolution IV design was compared to the best predicted experiment according to the model’s ME. The predicted best experiment showed a small (5.6%) but significant reduction on pCA production, confirming the importance of 2FI (Figure 2D, Sup. Table 4) .

### 3.3 Resolution V design: identification of relevant 2-factor interactions

Fractional factorial resolution V designs are required to identify ME and 2FI. When 11 factors are considered this results in 128 experiments. In order to decrease the number of experiments, factors with the highest impact on pCA production were fixed: only strains with high expression of PAL-C4H were considered and unbuffered media supplemented with Phe was used. This reduced the number of factors to 8 and the number of required experiments to 64. Considering the small variation of the resolution IV dataset, duplicates instead of triplicates were used during this round. The top producing conditions from the resolution IV experiments was included as control.

In the resolution V experiments, pCA production varied from 79.8 mg/l to 218.7 mg/l, improving the maximum production found in the first round by 38% (Figure 3A, Sup. Table 5). The use of strains with high expression of PAL-C4H in unbuffered media supplemented with Phe, increased the minimum production of pCA in this round by 62%, supporting the information provided by the resolution IV model.

**Figure 3:**
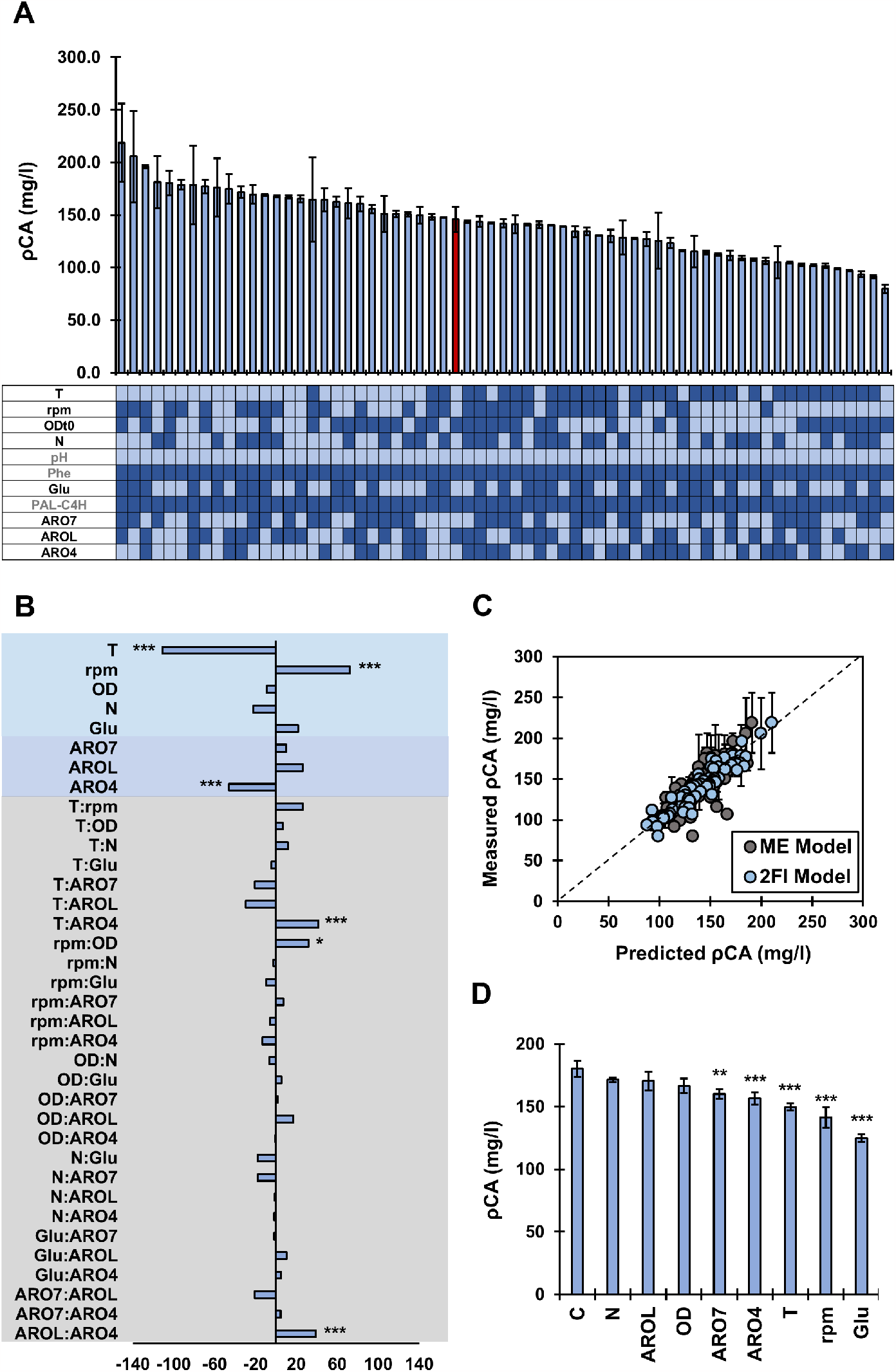
Resolution V Design **A**. Measured pCA production at different combinations of factors and levels. Light blue indicate low levels (-1) and dark blue indicates high level (1). The red bar is the best performing experiment from the resolution IV round. **B**. Coefficients of the 2FI model. *** indicates corrected p-value ≤ 0.001, ** ≤ 0.01 and * ≤ 0.05. **C**. Fit of models including main effects (ME) or main effects and two-factor interactions (ME-2FI model) to experimental data. **D** pCA production in validation experiment.

Experimental data was used to train new linear models based on ME or ME and 2FI. The model trained with ME showed a coefficient of determination of 0.55 that increased to 0.74 when 2FI were considered, highlighting the relevance of 2FI to explain pCA production (Figure 3C).

Temperature and agitation were identified as significant process related factors, so low T and high rpm improve pCA production (Figure 3 B). ARO4 was the only significant genetic factor, and, in contrast to other reports, low expression of this gene positively affected pCA titers (Figure 3 B) [10, 12]. Moreover, three significant positive 2FI were found: T:ARO4, rpm:OD and AROL:ARO4 (Figure 3 B).

In order to find the optimal strain and conditions for pCA production, the model including ME and 2FI was used to predict pCA titers for all strains in all possible media conditions. The use of a strain with high expression of PAL-C4H, ARO7 and AROL and lower expression of ARO4 in a media supplemented with urea, Phe and Glu incubated at 20°C and 250 rpm with an initial OD of 0.3 was predicted to optimize pCA production. These conditions were met by the top producing experiment measured in the resolution V round. To avoid the bias towards performed experiments during the estimation of model parameters, a new model was trained excluding data from the top producing experiment. When pCA production was predicted, the excluded top producer experiment was also suggested as optimal. Moreover, we evaluated the effect of individually changing each factor to their sub-optimal level. As expected, these modifications decreased or did not affect pCA production (Figure 3D, Sup. Table 6). Changing the initial OD, the nitrogen source or the expression of AROL all factors with not significant ME did not significantly change the pCA produced. In contrast, modifying the expression of ARO4, T and rpm, factors with significant ME, negatively impacted pCA production. Although ME related to ARO7 and Glu were insignificant, reducing the expression of ARO7 and omitting Glu supplementation, negatively impacted pCA production, which could be explained by higher-order factor interactions not included in the model.

Interaction graphs were used to understand the relationship between factors involved in significant 2FI and pCA production. Figure 4A shows the interaction between a genetic factor, ARO4, and a process-related factor, temperature. While at 30°C expression of ARO4 does not affect production, a lower expression results in a higher titer at 20°. Genetic factors also interact among each other, as an imbalanced expression of low AROL and high ARO4 results in the lowest pCA titer (Figure 4B). Last, a significant interaction between rpm and OD was found since, although fast agitation is always preferred, it has a higher impact in clotures with high initial cell density (Figure 4C).

**Figure 4:**
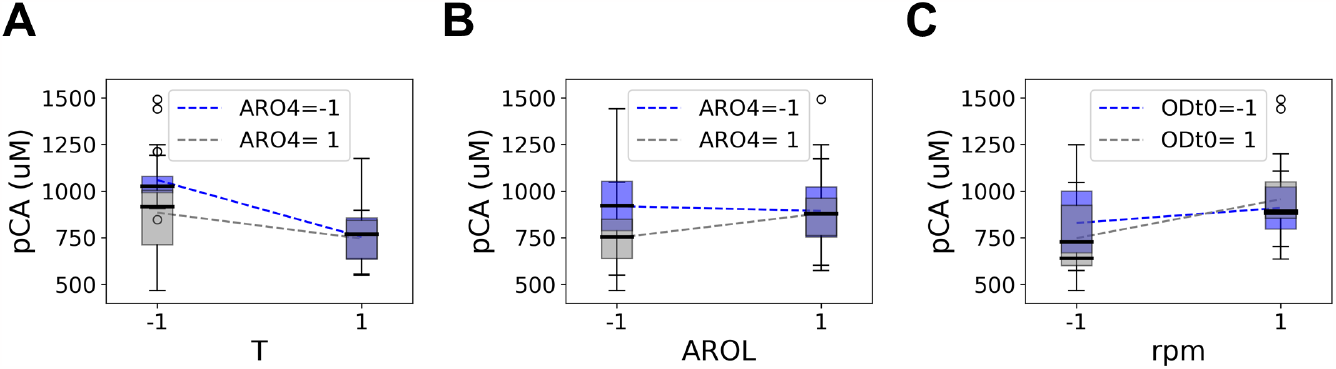
Interaction plots of significant 2-factor interactions. See Table 2 to identify levels corresponding to -1 and 1. T, temperature; rpm, agitation speed; ODt0, initial cell density.

## 4 Discussion

DoE has been commonly applied to process optimization [6, 23–25, 28, 29, 31] and strain design [5, 32, 33]. However, only a few studies consider simultaneous optimization of genetic and environmental factors [7, 8]. Importantly, these studies showed that the interplay between both types of factors must be taken into account simultaneously for optimization of microbial conversion processes. Our work underscores and strengthens these findings, using pCA production in *S. cerevisiae* as example. The importance of simultaneous process and strain optimization is highlighted by the T:ARO4 interaction (Figure 4A). If strain selection had been performed at *S. cerevisiae*’s standard growth temperature (30 °C), tuning the expression of ARO4 would have been considered irrelevant. However, given that the expression of ARO4 becomes important at lower temperatures, a sub-optimal strain could have been selected for subsequent process optimization.

The interplay between the strain performance and the bio-process design is especially important when moving from laboratory-scale to large-scale processes. This step-wise endeavour is time-consuming, labour-intensive and expensive. It thus benefits from scale-down experimentation [34]. The central paradigm of scaling-down states that scale-up will succeed when changes in the cellular environment caused by changes in scale do not influence cell behaviour [35]. Here we show how DoE can identify genetic and process parameters with significant influence on production that should be the focus of the down/up-scaling plan. We used DoE to understand the effect of 7 process-related factors (T, rpm, OD, N, pH, Phe and Glu), 4 genetic factors (PAL-C4H, ARO7, AROL and ARO4) and their interactions on pCA production. Considering two levels per factor, 2048 experiments would be required to find all possible interactions between factors and ensure the identification of the best production conditions. Instead, we performed 92 experiments (4.5% of the total) divided in two consecutive rounds. The first round identified the expression of PAL-C4H, the addition of Phe and the use of unbuffered media as key variables to ensure high pCA titers. These findings were taken up in a second experimental round for which the optimal pCA production conditions were found: incubation at low temperature and high agitation of a strain with low expression of ARO4. A 38% increased pCA production was obtained in the resolution V round compared to the best experiment in the resolution IV design. Moreover, a 168-fold improvement was measured between the worst and best performed experiments.

As indicated, only fourteen of the sixteen strains required for the resolution IV experiments were constructed, so only 28 out of the 32 designed experiments could be performed. Although resolution IV designs with 32 experiment allow the estimation of ME for up to 16 factors, we considered the impact of 11 factors on production. This granted some redundancy in the design and allowed the estimation of all ME with 28 out of the 32 designed experiments. However, if more factors had been considered, the construction of all the required strains would have been necessary for the estimation of ME. This limitation can be solved with the machine learning (ML) analysis of strain libraries generated using one-pot random transformation [15, 36]. Still, although ML can identify significant factors with an impact on production, quantifying the impact of interactions between factors, critical for bio-process optimization, is not trivial [15]. Moreover, the randomization of strains and process conditions, even when mini-bioreactor systems are used [37, 38], increases the complexity of creating suitable datasets for ML.

Summarizing, through a systematic evaluation of 11 factors, including genetic modifications and process parameters, we uncovered some of the interplay between genetic and environmental factors in pCA production. Moreover, we demonstrate the power of DoE to provide insights into factor effects and interactions for process optimization. By leveraging the strengths of experimental design and statistical analysis, we provide a framework to find key factors that impact bio-process performance that could be systematically applied to guide strain design as well as scale-up/down efforts.

## Acknowledgment

This project was founded by the Netherlands Organization for Scientific Research (NWO; project number GSGT.2019.008) and the European Union’s Horizon 2020 research and innovation program under grant agreement 814408 (Shikifactory100).

